# Cortico–thalamo–cerebellar circuit disruptions underlying cognitive deficits in schizophrenia: a multi-cohort multidomain fusion study

**DOI:** 10.1101/2025.06.02.657541

**Authors:** KuaiKuai Duan, Rogers F. Silva, Md Abdur Rahaman, Zening Fu, Jingyu Liu, Peter Kochunov, Theo G.M. van Erp, Sarah Shultz, Vince D. Calhoun

## Abstract

Disrupted large-scale brain network interactions are a hallmark of schizophrenia, yet the circuit-level mechanisms underlying cognitive deficits remain incompletely understood. In particular, cortico–thalamo–cerebellar circuitry has been implicated in “cognitive dysmetria,” but its functional organization across subdomains has not been fully characterized. Here, we apply a flexible multimodal data fusion framework (aNy-way independent component analysis) to integrate 4D domain-specific resting-state fMRI data from the cortex, thalamus, and cerebellum. Using a large discovery cohort (FBIRN, N=311) and two independent replication datasets (COBRE, N=148; MPRC, N=364), we identify a robust cortico–thalamo–cerebellar circuit involving higher-order thalamic nuclei, visual cortex, default mode network, and cerebellar posterior regions. This circuit shows altered connectivity in schizophrenia, characterized by reduced anticorrelation between thalamus and cortex, and is consistently associated with cognitive impairments, including processing speed, working memory, and reasoning deficits across cohorts. Importantly, these findings are largely replicated across independent cohorts. Together, these results provide data-driven evidence for a cortico–thalamo–cerebellar mechanism underlying cognitive dysfunction in schizophrenia, supporting the cognitive dysmetria hypothesis. This work also demonstrates the utility of aNy-way independent component analysis for identifying clinically relevant brain circuits in psychiatric disorders.

**Highlights:**

- Identified cortico–thalamo–cerebellar circuits in schizophrenia
- Altered thalamocortical interactions differentiated schizophrenia
- Circuit alterations were associated with cognitive impairment
- Findings are replicated across three independent multisite datasets
- Multidomain fusion enabled subdomain-specific circuit characterization

## 1. Introduction

Schizophrenia (SZ) is characterized by widespread disruptions in large-scale brain network interactions, particularly affecting cognitive functions such as working memory, attention, and executive functioning (Calhoun et al., 2009; Dong et al., 2018; Menon et al., 2023). A prominent hypothesis, “cognitive dysmetria”, suggests that dysfunction in cortico–thalamo–cerebellar circuitry may underlie these impairments (Andreasen et al., 1998). The thalamus plays a central role in integrating sensory information and coordinating communication between cortical regions (Hwang et al., 2017). Impairments in thalamic structure (Pergola et al., 2015) and thalamocortical connectivity (Anticevic et al., 2014; Avram et al., 2018; Chen et al., 2019; Gong et al., 2019; Sheffield et al., 2020; Yao et al., 2020) have been extensively documented in SZ patients. Similarly, cerebellum, contributes to a wide range of cognitive processes, including motor learning, working memory, emotion, and language (Ren et al., 2019), and exhibits both anatomical and functional alterations in SZ (Andreasen and Pierson, 2008; Ding et al., 2019; Li et al., 2022). Aberrant thalamus-cerebellar connectivity (Ha et al., 2023), altered cerebro-cerebellar connections (Collin et al., 2012; Kang et al., 2024; Kim et al., 2020), and disrupted cortico-thalamo-cerebellar connectivity (Ha et al., 2023) have also been reported in SZ. However, these studies have largely employed region of interest (ROI)-based approaches relying on anatomical definitions of fixed, binary, and nonoverlapping ROIs without leveraging functional and connectivity information, which may not fully capture functionally relevant subregions or their interactions.

Parallel to these findings, there has been increasing interest in characterizing interactions across multiple brain domains, including the thalamus, cerebellum, and cortical regions, using data-driven approaches (Chen et al., 2019; Collin et al., 2012; Ferri et al., 2018; Kim et al., 2020). Traditional data-driven functional connectivity approaches, e.g., group independent component analysis (GICA) (Calhoun et al., 2001), typically decompose whole-brain fMRI data using a single model order (Calhoun and Adali, 2012). While this may not be optimal as different brain domains have different spatial organizations (Bijsterbosch et al., 2018; Diedrichsen, 2006; Iglesias et al., 2018; Tzourio-Mazoyer et al., 2002) and subdivisions of specific domains (e.g., thalamus) have distinct connectivity with other domains (e.g., cortex and cerebellum) (Giraldo-Chica and Woodward, 2017; Gong et al., 2019; Ha et al., 2023; Yuan et al., 2016; Zhang et al., 2021). Recent studies employing domain-wise brain decomposition have revealed crucial insights on brain functionalities and enhanced sensitivity in detecting subdomain networks potentially overlooked in whole-brain connectivity studies (Dobromyslin et al., 2012; Hwang et al., 2017). Meanwhile, data-driven functional parcellations of specific regions or domains, such as the thalamus (Zhang and Li, 2017) and cerebellum (Dobromyslin et al., 2012; Ren et al., 2019), have revealed informative details at higher spatial scales and revealed distinct connectivity with other domains/networks (Giraldo-Chica and Woodward, 2017; Gong et al., 2019; Ha et al., 2023; Yuan et al., 2016).

To date, no study has comprehensively investigated cortico–thalamo–cerebellar circuit organization using a data-driven fusion framework that allows for domain-specific resolution and flexible modeling of inter-domain relationships. A flexible multimodal fusion framework, aNy-way independent component analysis (aNy-way ICA), has been previously introduced to enable integration of multiple modalities or domains with distinct model orders for each modality/domain (Duan et al., 2020). The flexibility of aNy-way ICA makes it better suited to capture the complex and heterogeneous neurobiology of SZ. When integrating multidomain data, aNy-way ICA simultaneously conducts domain-specific granular parcellation and optimizes connections between subdivisions of different domains. While the aNy-way ICA framework has been extensively validated in simulation studies (Duan et al., 2020), its utility for identifying clinically meaningful brain circuits in psychiatric disorders remains unclear.

In this study, we apply aNy-way ICA to integrate 4D multidomain resting-state fMRI data to comprehensively identify cortico–thalamo–cerebellar functional circuits in schizophrenia. Using a large discovery cohort of 311 subjects (151 with SZ) from the Function Biomedical Informatics Research Network (FBIRN) (Keator et al., 2016) and two independent replication datasets, including 364 subjects from the Maryland Psychiatric Research Center (MPRC) cohort (Adhikari et al., 2019) ), and 148 individuals from the Center for Biomedical Research Excellence (COBRE) cohort (Aine et al., 2017), we aim to (1) identify subdomain-specific cortico–thalamo–cerebellar linkages, (2) examine their ability to differentiate individuals with SZ from healthy controls, and (3) evaluate their associations with cognitive performance and symptom severity. By leveraging a data-driven approach with flexible domain-specific modeling, this study seeks to provide new insights into circuit-level mechanisms underlying cognitive dysfunction in SZ.

## 2. Materials and Method

### 2.1. Samples

Resting state fMRI data from three independent multicenter studies, including FBIRN (Keator et al., 2016), COBRE (Aine et al., 2017), and MPRC (Adhikari et al., 2019), were analyzed to identify and replicate cortico-thalamo-cerebellar functional circuits and whole-brain functional network connectivity (FNC) associated with SZ. These datasets included individuals with SZ and healthy controls (HCs) and enabled evaluation of the robustness and reproducibility of the identified circuit-level findings across independent cohorts. Each study was approved by the institutional review board and all participants provided written informed consent.

The discovery sample was composed of 311 individuals (age: 37.88±11.26 years) recruited from the FBIRN project that also passed quality control (QC, see details later) (Keator et al., 2016), including 151 individuals with SZ (36 female) and 160 HCs (45 female). All participants are adults (18-62 years old), and their cognitive performance was measured by the Computerized Multiphasic Interactive Neurocognitive System (CMINDS) (van Erp et al., 2015). Diagnosis of SZ was determined using DSM-IV (American Psychiatric, 1994). Table 1 summarizes the demographic and cognitive scores of included FBIRN subjects. Age and sex were matched between individuals with SZ and HCs (Table 1).

**Table 1.** Demographic Information and Cognitive Scores for SZ and HC Individuals in the FBIRN Cohort.

| Characteristics | HC | SZ | p value (SZ vs. HC) |
| --- | --- | --- | --- |
| Number | 160 | 151 |  |
| Sex (M/F) | 115/45 | 115/36 | 0.39<sup>a</sup> |
| Age (years) | 37.04 (10.86) | 38.77 (11.63) | 0.18<sup>b</sup> |
| **CMINDS** |  |  |  |
| Composite score | 0.04 (0.95) | -1.59 (1.22) | $3.59 \times 10^{-28}$ b |
| Speed of processing | 0.06 (1.02) | -1.34 (1.08) | $7.28 \times 10^{-24}$ b |
| Attention/vigilance | 0.03 (0.99) | -1.36 (1.40) | $6.15 \times 10^{-19}$ b |
| Working memory | 0.05 (0.97) | -1.16 (1.08) | $6.71 \times 10^{-20}$ b |
| Verbal learning | 0.04 (1.00) | -1.38 (1.19) | $5.50 \times 10^{-23}$ b |
| Visual learning | 0.03 (0.99) | -1.09 (1.14) | $3.97 \times 10^{-16}$ b |
| Reasoning/problem solving | 0.04 (0.97) | -0.85 (1.22) | $1.21 \times 10^{-10}$ b |

<sup>a</sup>Pearson's chi-squared test.
<sup>b</sup>Welch's two sample t test.

**Table 2.** Demographic Information and Cognitive Scores for SZ and HC Individuals in the COBRE Cohort. Characteristics HC SZ p value (SZ vs. HC)

| Characteristics | HC | SZ | p value (SZ vs. HC) |
| --- | --- | --- | --- |
| Number | 83 | 65 |  |
| Sex (M/F) | 61/22 | 54/11 | 0.16<sup>a</sup> |
| Age (years) | 38.35 (11.84) | 37.26 (13.94) | 0.18<sup>b</sup> |
| **MCCB** |  |  |  |
| Composite score | 49.92 (8.69) | 31.68 (13.62) | $7.89 \times 10^{-17}$ b |
| Speed of processing | 53.81 (8.62) | 34.97 (11.77) | $1.65 \times 10^{-21}$ b |
| Attention/vigilance | 49.99 (9.21) | 37.38 (13.29) | $5.78 \times 10^{-10}$ b |
| Working memory | 48.92 (10.67) | 39.28 (12.62) | $1.41 \times 10^{-6}$ b |
| Verbal learning | 45.53 (8.32) | 38.66 (8.78) | $2.95 \times 10^{-6}$ b |
| Visual learning | 46.02 (10.71) | 35.37 (12.70) | $1.41 \times 10^{-7}$ b |
| Reasoning/problem solving | 55.74 (8.11) | 44.02 (10.93) | $1.13 \times 10^{-11}$ b |

<sup>a</sup>Pearson's chi-squared test.
<sup>b</sup>Welch's two sample t test.

**Table 3.** Demographic Information of SZ and HC Individuals in the MPRC Cohort.

| Demographics | HC | SZ | p value (SZ vs. HC) |
| --- | --- | --- | --- |
| Number | 229 | 135 |  |
| Sex (M/F) | 143/86 | 49/86 | $1.39 \times 10^{-6}$<sup>a</sup> |
| Age (years) | 40.52 (15.22) | 38.79 (13.82) | 0.28<sup>b</sup> |

<sup>a</sup>Pearson's chi-squared test.
<sup>b</sup>Welch's two sample t test.

### 2.2. fMRI data acquisition and preprocessing

The acquisition information of each fMRI dataset can be found in the supplementary information. All fMRI data were preprocessed using statistical parametric mapping (SPM12, http://www.fil.ion.ucl.ac.uk/spm/) following the NeuroMark fully automated spatially constrained ICA pipeline (Du et al., 2020; Fu et al., 2021) implemented in the GIFT software (http://trendscenter.org/software/gift). Specifically, subject head motion was corrected using rigid body motion correction, followed by slice-timing correction to adjust timing difference in slice acquisition. The slice-timing corrected fMRI data were further normalized to the standard Montreal Neurological Institute (MNI) space using an echo planar imaging (EPI) template (Calhoun et al., 2017). Normalized fMRI data were resampled to 3 × 3 × 3 mm^3^ (for extraction of thalamus and cerebellum) and 6 × 6 × 6 mm^3^ (for extraction of regions excluding thalamus and cerebellum, see details later) isotropic voxels separately and further spatially smoothed using a Gaussian kernel with a full width at half maximum (FWHM) of 6 mm.

NeuroMark QC (Chen et al., 2022; Du et al., 2020; Fu et al., 2021; Thapaliya et al., 2025) was applied to all preprocessed scans. We excluded (1) subjects with head motions of any frames greater than 3 mm translation or exceeding 3° rotation (compared to the first frame after discarding the first five frames to account for signal equilibrium and participants’ adaptation to the scanner’s noise); (2) subjects with number of time points less than 120; (3) subjects with poor normalization quality.

### 2.3. fMRI feature extraction in the thalamus, cerebellum, and cortex

Thalamus, cerebellum and cortex regions (i.e., all other regions excluding thalamus and cerebellum, referred it as cortex since they are primarily cortical regions) were defined and extracted from preprocessed fMRI data using the automated anatomical labeling (AAL) atlas (Tzourio-Mazoyer et al., 2002). To make feature dimensions of subdomains (thalamus, cerebellum, cortex) compatible and speed up the convergence of domain-wise fMRI data fusion, thalamus and cerebellum were extracted from fMRI data with 3 × 3 × 3 mm^3^ resolution, yielding 609 voxels for thalamus and 7217 voxels for cerebellum. Cortical regions were extracted from fMRI data with 6 × 6 × 6 mm^3^ resolution, yielding 5916 voxels.

### 2.4. Cortico–thalamo–cerebellar subdomain fusion using aNy-way ICA

In this study, we apply a previously developed multimodal fusion framework, aNy-way ICA (Duan et al., 2020), to integrate multidomain resting-state fMRI data. Below, we provide a brief overview of the aNy-way ICA framework and focus on its application to cortico–thalamo–cerebellar circuit analysis in schizophrenia. Detailed methodological derivations and simulation validations have been described in (Duan et al., 2020).

#### 2.4.1. aNy-way ICA

The aNy-way ICA (Duan et al., 2020) framework provides a flexible approach for integrating multidomain neuroimaging data and enabling identification of linked functional circuits across brain regions. aNy-way ICA aims to maximize statistical independence within each domain while minimizing mutual information among subspace component vectors (SCVs, SCVs represent linked components across domains). Fig.1 shows the conceptual overview of the aNy-way ICA framework applied to multidomain neuroimaging data. Without loss of generality, we consider three domains: cortex, thalamus, and cerebellum, each with 3, 4, and 5 sources, respectively. For each domain *m*, the observed data 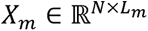 is decomposed into a source matrix *s* ϵ 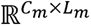 and a loading matrix 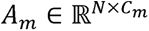, such that:

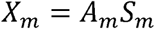

where *N* = number of subjects x total number of time points, *L_m_* is the feature dimensionality, and *c_m_* is the number of components for domain *m*. Importantly, aNy-way ICA allows different numbers of components across domains.

**Fig. 1.**
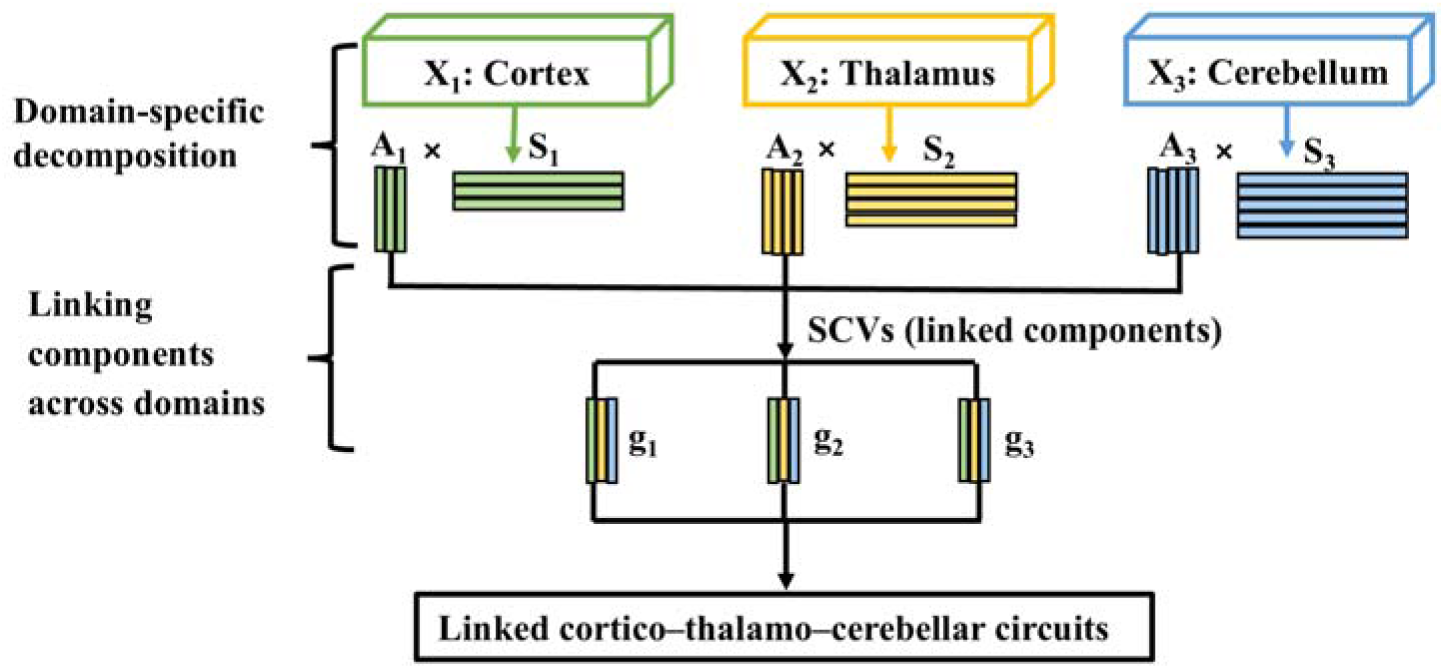
Conceptual overview of multidomain fusion for identifying cortico–thalamo–cerebellar circuits using aNy-way ICA. Data from multiple brain domains (cortex, thalamus, and cerebellum) are decomposed into domain-specific components (**S**_m_, m = 1,2,3) and associated subject loadings (**A**_ill_,m = 1,2,3). Components that have shared patterns of variation across domains are then aligned to form subspace component vectors (SCVs, **g**_k_, k = 1,2,3). Each SCV represents a cortico–thalamo–cerebellar circuit, defined as a set of functionally coupled patterns spanning cortical, thalamic, and cerebellar subregions.

To model cross-domain relationships, corresponding loadings across domains are grouped into subspace component vectors (SCVs), representing linked components across domains. The total number of SCVs is determined by the minimum number of components across domains. These SCVs capture shared variability across domains while allowing domain-specific structure to be preserved.

The aNy-way ICA framework jointly optimizes within-domain independence and between-domain linkage using the following cost function:

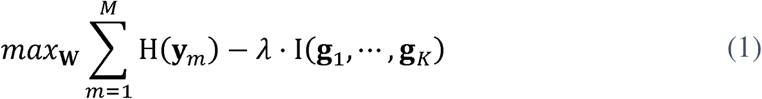

where H(*y_m_*) represents the entropy of the nonlinearly transformed sources for domain m, promoting independence. *M* is the total number of domains. I(**g**_1_,…, **g***_K_*) denotes the mutual information among SCVs, encouraging linkages across domains. The parameter ,*λ* balances these two objectives. Entropy is defined following standard infomax ICA (Bell and Sejnowski, 1995) and mutual information is defined following standard independent vector analysis (Anderson et al., 2012; Bell and Sejnowski, 1995). SCVs are modeled using a multivariate Gaussian assumption (IVA-G), allowing second-order statistical dependencies across modalities to be captured (Anderson et al., 2012). Optimization details are described in (Duan et al., 2020).

#### 2.4.2. Identify cortico–thalamo–cerebellar functional linkages using aNy-way ICA

For each subdomain (thalamus, cerebellum, cortex) fMRI data, time courses were concatenated along subject direction to form a two-dimensional feature matrix with dimensions *N*x *L* (*N* = total number of subjects × number of time points; *L* = number of voxels). Subject-wise mean removal was then applied, and the resulting data were input into aNy-way for domain-wise data fusion. aNy-way ICA simultaneously maximizes the independence of components within each subdomain while maximizing the connectivity between these subdomains. The component number was estimated separately for each subdomain based on the order selection method proposed in (Li et al., 2007). Assuming the minimum component number across the thalamus, cerebellum, and cortex is *K*, aNy-way ICA can effectively optimize *K* cortico-thalamo-cerebellar functional linkages (i.e., triplets or SCVs).

After aNy-way ICA converged, the sources (representing spatial maps of intrinsic connectivity networks (ICNs)) and their corresponding loadings (representing time courses (TC)) were obtained for the thalamus, cerebellum, and cortex, separately. A series of postprocessing steps were conducted on the TCs to eliminate noise sources, including (1) removal of linear, quadratic, and cubic trends; (2) adjusting motion effect using a multiple regression model with six realignment parameters and their temporal derivatives as independent variables; (3) de-spiking to detect and exclude outliers; and (4) band-pass filtering with [0.01–0.15] Hz.

Beyond identifying linked subdomain SCVs, aNy-way ICA decompositions also enable the computation of whole-brain FNC to further investigate their group differences (SZ vs. HC). Both *K* cortico-thalamo-cerebellar functional linkages and whole-brain FNC were derived from the postprocessed TCs. Specifically, *K* cortico-thalamo-cerebellar triplets (i.e., SCVs) were formed by concatenating the postprocessed TCs of the first *K* components from the thalamus, cerebellum, and cortex subdomains. Cross-correlation of these *K* triplets was computed using

Pearson correlation and then Fisher-transformed to quantify cortico-thalamo-cerebellar functional linkages. Whole-brain FNC was computed in a similar manner, with cross-correlation estimated using all components from the thalamus, cerebellum, and cortex subdomains.

Multiple comparison was corrected over whole-brain FNCs using a false discovery rate (FDR) of *p* < 0.05 (‘mafdr’ command in Matlab 2023a) (Duan et al., 2021b; Storey and Tibshirani, 2003) to take dependencies of connections within SCVs into account. SCVs were considered significantly interlinked when all their pair-wise connection were whole-brain wide significant (i.e., passed FDR at *p* < 0.05 correction over whole-brain FNCs).

### 2.5. Statistical inference

For each significantly interlinked SCV, we computed pair-wise subdomain connection for each participant and examined their ability for discriminating SZ from controls using a N-way analysis of variance (ANOVA) model:

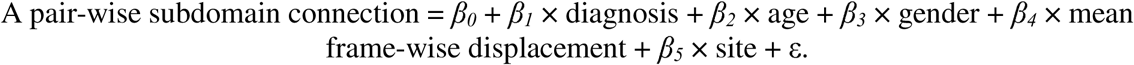

where each pair-wise subdomain connection is treated as the response variable and diagnosis is the predictor. Age, sex, mean frame-wise displacement (FD) and site are covariates of no interest. FDR at *p* < 0.05 was applied to correct for comparisons from all pair-wise subdomain connections tested. SZ vs. HC difference of the whole-brain FNC was also examined using the abovementioned N-way ANOVA model, and multiple comparison was corrected over all whole-brain FNCs at FDR *p* < 0.05. We further tested the associations between pair-wise subdomain connections and cognitive variables listed in Tables I-II using a N-way ANOVA model:

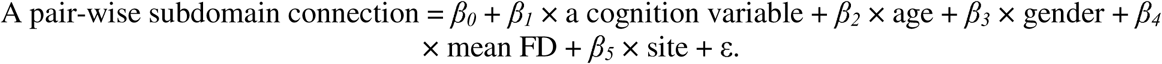

Similarly, each pair-wise subdomain connection is treated as the response variable and age, sex, mean FD and site are covariates of no interest. Cognitive variables are the predictors. FDR at *p* < 0.05 was applied to correct for comparisons from all pair-wise subdomain connections tested.

### 2.6. Replicability of subdomain connections

We assessed the replicability of the identified subdomain linkages, their group differences (SZ vs. HC) and associations with cognition, as well as whole-brain FNC group differences in COBRE and MPRC datasets. For each replication dataset, we applied the same preprocessing steps as in the discovery (FBIRN) dataset and projected the decomposed subdomain ICNs from FBIRN onto COBRE and MPRC data to extract the corresponding TCs (Duan et al., 2023; Duan et al., 2021a). The resulting TCs underwent the same post-processing steps, including detrending, motion correction, de-spiking, and band-pass filtering. Since cognitive performance data included in FBIRN were not available in the MPRC dataset but were available in comparable formats for the COBRE dataset, we specifically examined the replicability of associations between subdomain connections and cognition in the COBRE dataset.

Consistent with the FBIRN dataset, we used Pearson correlation tests to evaluate subdomain linkages and applied N-way ANOVA models to assess group difference (SZ vs. HC) and associations with cognition for each pair-wise subdomain connection in the COBRE and MPRC datasets. A pair-wise subdomain linkage was considered replicated if its p-value was less than 0.05 and it exhibited the same directionality as in the FBIRN dataset. The group difference (SZ vs. HC) for a pairwise subdomain connection was considered replicated if (1) the connection was replicated; (2) the p-value for SZ vs. HC difference was below 0.05; and (3) the directionality aligned with the FBIRN findings. A similar criterion was applied to assess the replicability of associations between pair-wise subdomain connections and cognition in the COBRE dataset.

## 3. Results

We estimated 7 components for the thalamus, 11 components for the cerebellum, and 29 components for the cortex using the method in (Li et al., 2007). Thus, the total number of SCV is 7 (i.e., the minimum component number among the three subdomains). Among the identified seven subdomain SCVs, one SCV (Fig. 2, SCV 6) was significantly interlinked after FDR at *p* < 0.05 correction. The thalamus component (blue color in Fig. 2 (a)) was positively and significantly connected with the cerebellum component (green color in Fig. 2 (a), *r* = 0.16, *p* = 6.19×10^-3^, Fig. 2 (b)). The cortex component (hot color in Fig. 2 (a)) was significantly anti-correlated with the thalamus (*r* = −0.15, *p* = 1.01×10^-2^, Fig. 2 (c)) and the cerebellum component (*r* = −0.20, *p* = 5.41×10^-4^, Fig. 2 (d)). Supplementary Fig. 1 plots the identified thalamus, cerebellum, and cortex components overlaid to their corresponding masks. The thalamus component highlighted medial and posterior thalamus (e.g., the mediodorsal and the pulvinar nuclei). The cerebellum component highlighted Crus I and declive, and the cortex component highlighted cuneus, precuneus, lingual, and posterior cingulate cortex.

**Fig. 2.**
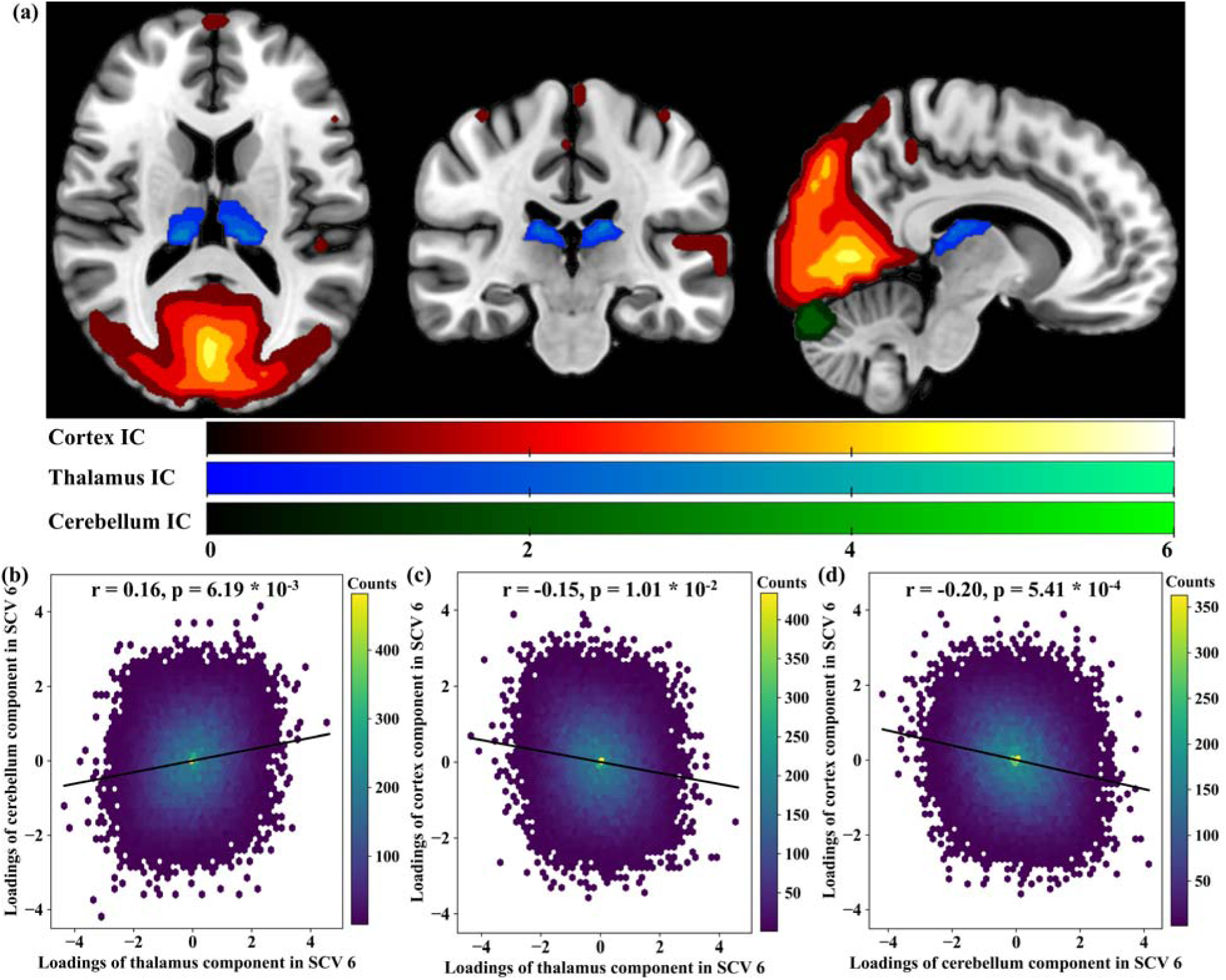
**The identified cortico-thalamo-cerebellar SCV and their linkages in the FBIRN (discovery) dataset**. (a) The Z-scored spatial map of the identified cortico-thalamo-cerebellar SCV and (b-d) corresponding pairwise connections: (b) thalamus-cerebellum, (c) thalamus-cortex, (d) cerebellum-cortex.

The connection between thalamus and cortex in SCV 6 showed significant difference between SZ and HC (Fig. 3(a), *p* = 3.77×10^-7^, Cohen’s d (referred as d hereafter) = 0.62), where SZ patients had weaker anti-correlation (i.e., hyperconnectivity) compared to HCs. Stronger thalamus-cortex connection in SCV 6 was significantly associated with lower CMINDS composite score (Fig. 3(b), *r* = −0.25, *p* = 1.87×10^-4^), slower processing speed (Fig. 3(c), *r* = - 0.18, *p* = 7.02×10^-3^), and poorer reasoning/problem solving skills (Fig. 3(d), *r* = −0.17, *p* = 8.09×10^-3^), poorer verbal learning ability (Fig. 3(e), *r* = −0.17, *p* = 7.48×10^-3^), poorer attention/vigilance (Fig. 3(f), *r* = −0.22, *p* = 1.15×10^-3^) and poorer working memory performance (Fig. 3(g), *r* = −0.18, *p* = 6.73×10^-3^). Stronger thalamus-cortex connection in SCV 6 showed nominal association with poorer visual learning abilities (Supplementary Fig. 2(a), *r* = −0.12, *p* = 0.04).

**Fig. 3.**
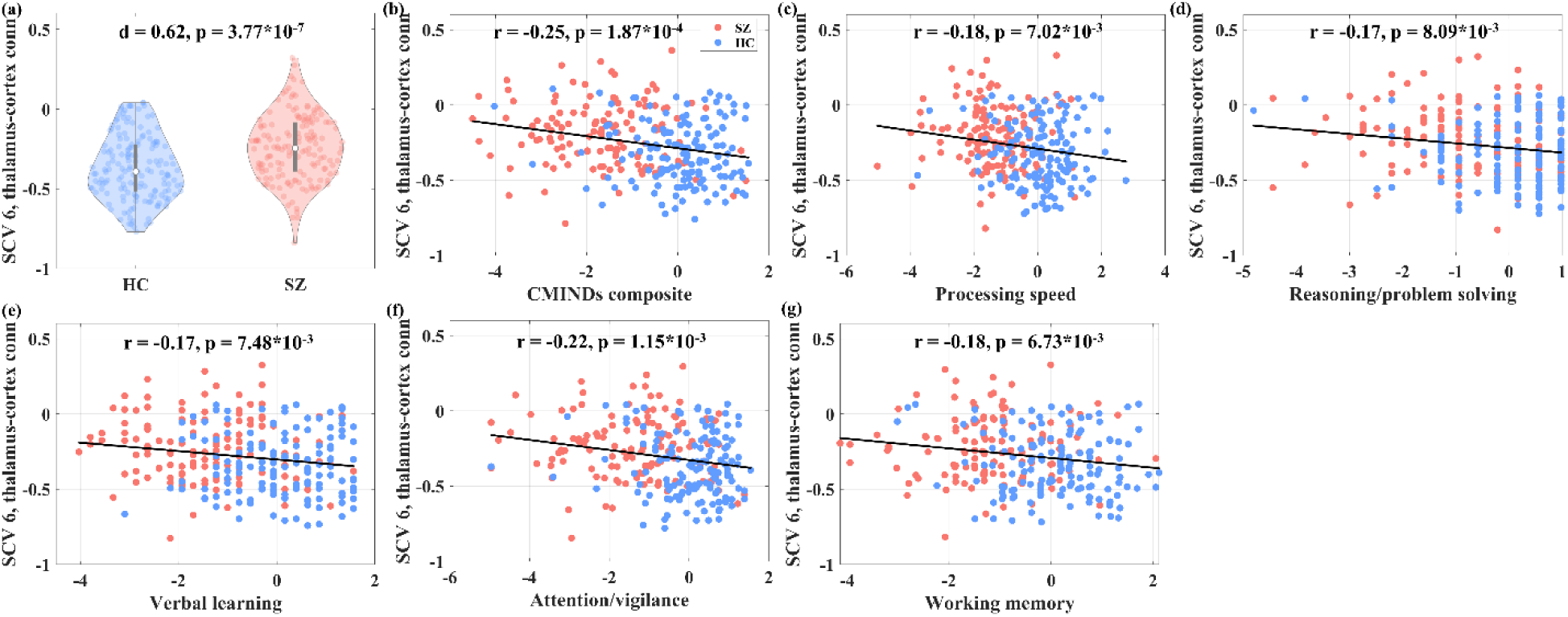
(a) Group difference and (b-g) associations with cognitive variables ((b) CMINDs composite, (c) processing speed, (d) reasoning/problem solving, (e) verbal learning, (f) attention/vigilance, (g) working memory) of the thalamus-cortex connection in SCV6 in the discovery dataset after adjusting effects from age, sex, site, and head motion. Red dots denote individuals with SZ and blue dots represent HCs.

In the whole-brain FNC analysis, 254 out of 1081 total connection pairs were significantly linked after FDR at *p* < 0.05 correction (see connectogram in Supplementary Fig. 3). Moreover, 47 out of the 254 significantly linked pairs showed significant SZ vs. HC difference after FDR at *p* < 0.05 correction. Fig. 4(a) plots the connectogram of 47 FNCs showing significant SZ vs. HC difference and the connectogram of corresponding SZ vs. HC difference (Cohen’s d value) is plotted in Fig. 4 (b).

**Fig. 4.**
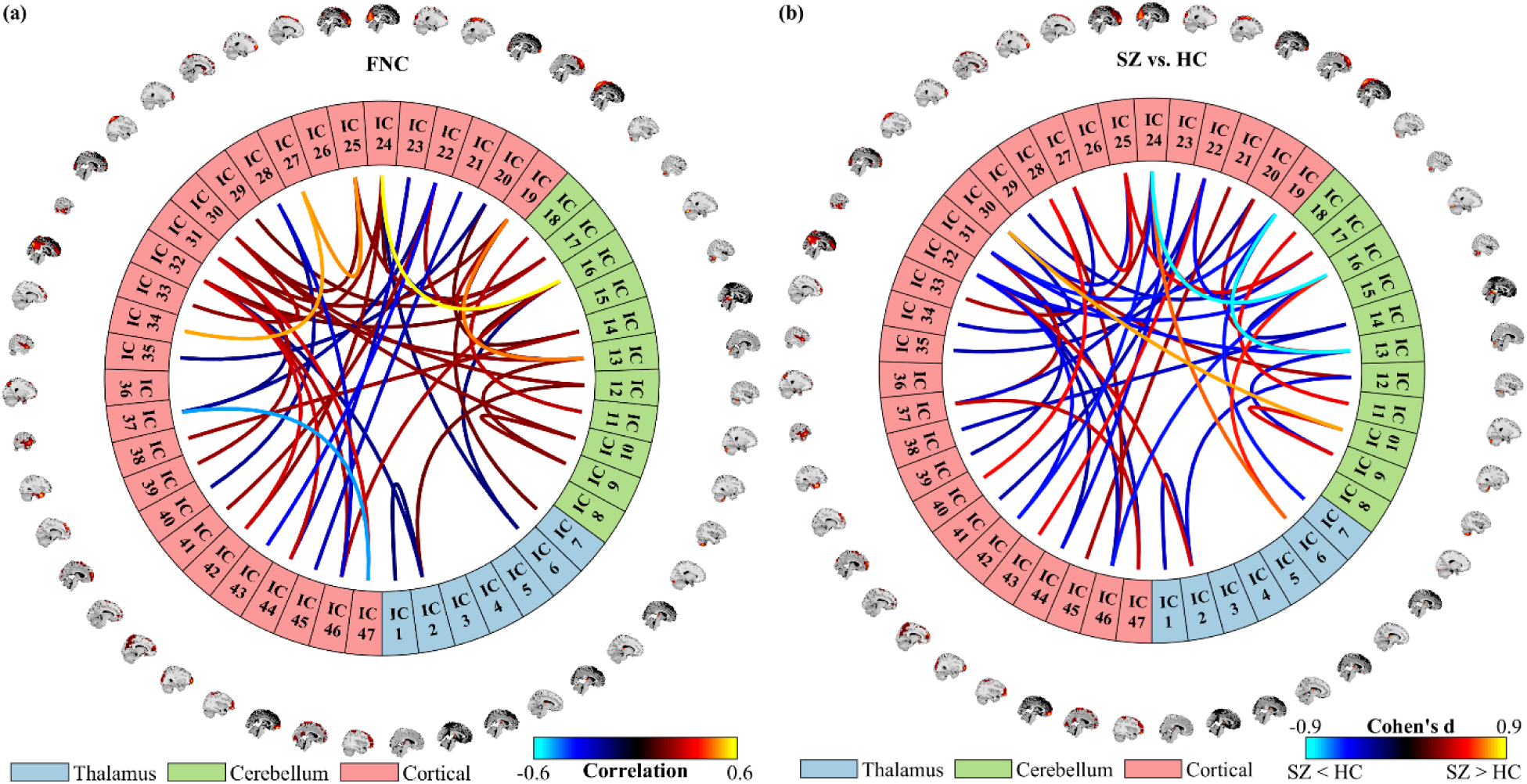
(a) The connectogram of significant FNCs (the color of the edge represents correlation values between subdomain components) and (b) the connectogram of FNCs showing significant SZ vs HC difference (the color of the edge represents Cohen’s d values of SZ vs. HC difference).

### 3.1. Replication in the COBRE dataset

The interconnections of the identified SCV 6 were replicated in the COBRE dataset (Fig. 5 (a)). The thalamus component had a significant and positive association with the cerebellum component (*r* = 0.19, *p* = 6.95×10^-3^, the left panel in Fig. 5 (a)). The cortex component showed significant anticorrelations with the thalamus (*r* = −0.22, *p* = 3.03×10^-3^, the middle panel in Fig. 5 (a)) and the cerebellum components (*r* = −0.21, *p* = 3.03×10^-3^, the right panel in Fig. 5 (a)).

**Fig. 5.**
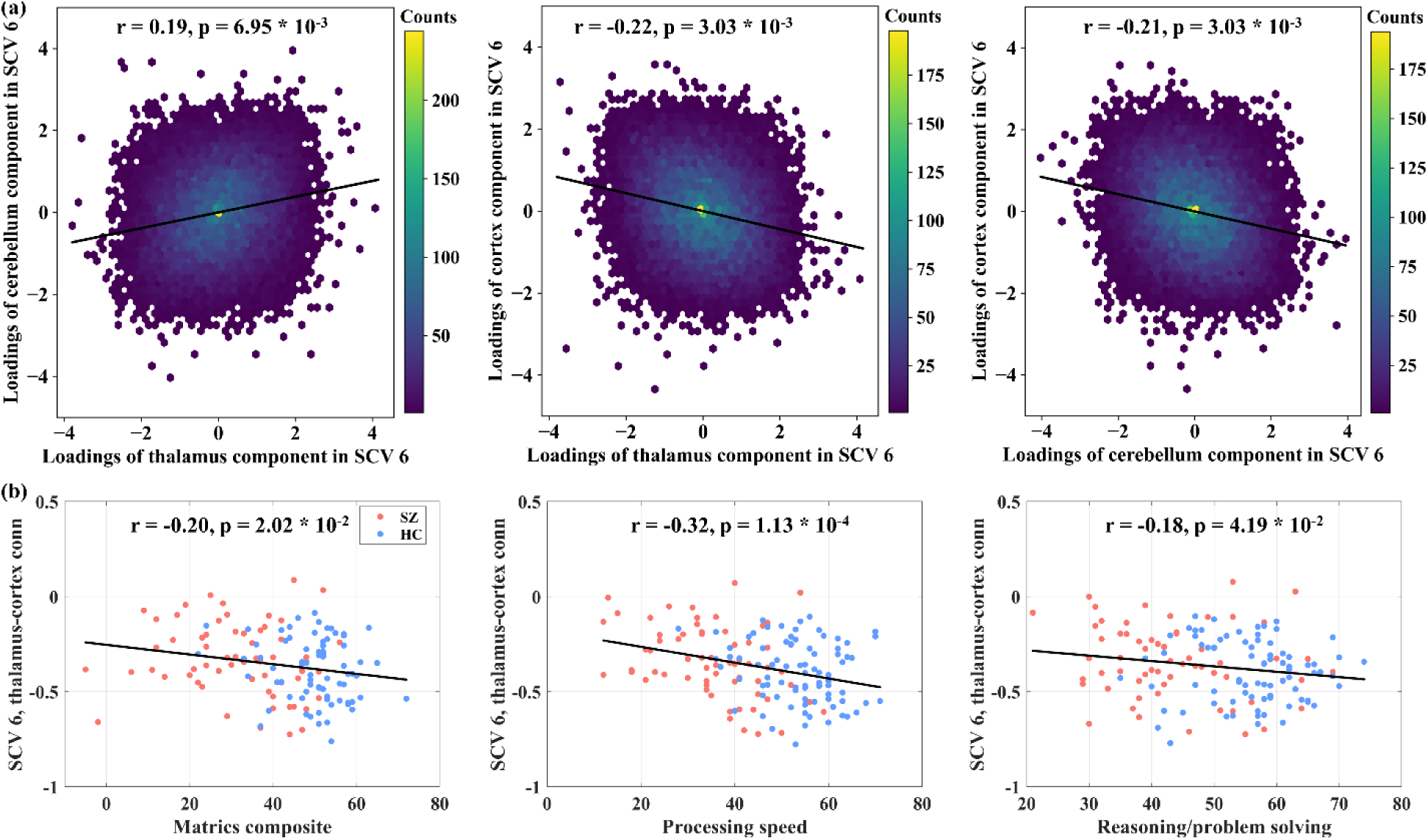
(a) The pair-wise connections of the identified SCV 6 and (b) the association between thalamus-cortex connection in SCV 6 and cognitive variables in the COBRE dataset after adjusting effects from age, sex, site and head motion. Red dots denote individuals with SZ and blue dots represent HCs.

In the COBRE dataset, the thalamus-cortex connection in SCV 6 was significantly and negatively associated with MATRICS composite score (Fig. 5 (b), *r* = −0.20, *p* = 2.02×10^-2^), processing speed (Fig. 5 (b), *r* = −0.32, *p* = 1.13×10^-4^), reasoning/problem solving skills (Fig. 5 (b), *r* = −0.18, *p* = 4.19×10^-2^), and visual learning abilities (Supplementary Fig. 2 (b), *r* = −0.21, *p* = 0.01). The thalamus-cortex connection in SCV 6 did not show significant difference between SZ and HCs (*p* = 0.09). In the whole-brain FNC analysis, 6 out of 47 FNCs showing significant SZ vs HC difference in the FBIRN (discovery) dataset were replicated in the COBRE dataset, and the connectogram of their Cohen’s d values for SZ vs. HC differences is plotted in Supplementary Fig. 4.

### 3.2. Replication in the MPRC dataset

Two out of the three interconnections of the identified SCV 6 were replicated in the MPRC dataset. The cortex component was significantly anti-correlated with the thalamus (*r* = −0.11, *p* = 0.03, Fig. 6 (a)) and the cerebellum components (*r* = −0.18, *p* = 6.71×10^-4^, Fig. 6 (b)). SZ patients also had significantly weaker anti-correlation between thalamus and cortex TCs in SCV 6 compared to HCs (d = 0.29, *p* = 9.70×10^-3^, Fig. 6 (c)). In the whole-brain FNC analysis, 11 out of 47 FNCs showing significant SZ vs HC difference in the FBIRN (discovery) dataset were replicated in the MPRC dataset, and the connectogram of their Cohen’s d values for SZ vs. HC differences is plotted in Fig. 6 (d).

**Fig. 6.**
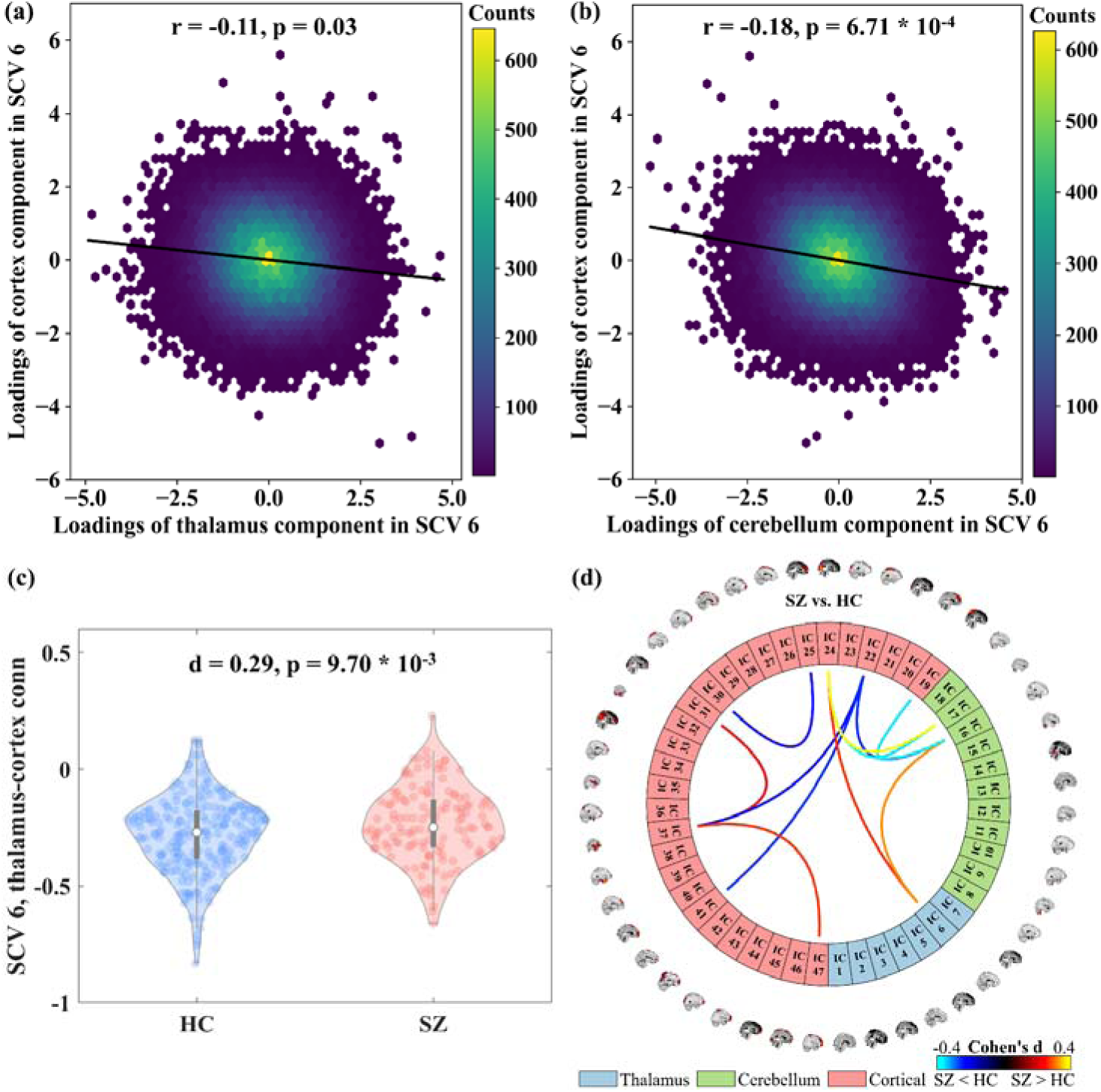
(a-b) The pair-wise connections of the identified SCV 6 and (c) group difference of the identified thalamus-cortex connection in SCV6 in the MPRC dataset after adjusting effects from age, sex, site and head motion. Red dots denote individuals with SZ and blue dots represent HCs. (d) The connectogram of FNCs replicated for SZ vs. HC difference in the MPRC dataset (the color of the edge represents Cohen’s d values of SZ vs. HC difference).

## 4. Conclusion

In this study, we identified and largely replicated a cortico-thalamo-cerebellar functional linkage (SCV 6) associated with cognitive deficits in schizophrenia across three independent datasets (FBIRN, COBRE and MPRC). This cortico-thalamo-cerebellar circuit highlights the functional connections among medial and posterior thalamus, cerebellar regions (Crus I and declive), and cortical regions in the visual network (e.g., cuneus and lingual gyrus) and default mode network (DMN, e.g., precuneus and posterior cingulate cortex). Notably, the anti-correlation between medial/posterior thalamus and cortex (mostly in the visual network and DMN) in SCV 6 was significantly reduced in individuals with SZ compared to HCs in both FBIRN and MPRC datasets. More importantly, stronger connectivity between medial/posterior thalamus and DMN/visual network was consistently related to more severe cognitive deficits in both FBIRN and COBRE datasets, including lower CMINDs/MATRICS composite score, slower processing speed, impaired visual learning abilities and reduced reasoning/problem-solving skills. Together, these findings provide evidence for a cortico–thalamo–cerebellar mechanism underlying cognitive dysfunction in schizophrenia.

The identified thalamus component highlights higher order thalamic nuclei, including the mediodorsal and the pulvinar nuclei, which are known to play key roles in integrating sensory information and supporting higher cognitive functions (Steiner et al., 2020; Tu et al., 2015; Yuan et al., 2016; Zhang et al., 2008). The mediodorsal and pulvinar have been consistently linked to DMN (Ha et al., 2023; Steiner et al., 2020; Yuan et al., 2016), and their connectivity was related to cognitive processes such as selective attention, processing speed, and cognitive flexibility (Steiner et al., 2020). Moreover, the connections between the visual cortex and the pulvinar nucleus have been well documented in humans (Cortes et al., 2024; Li et al., 2012; Sabatinelli et al., 2011; Steiner et al., 2020; Yuan et al., 2016; Zhang et al., 2008; Zuo et al., 2010), monkeys (Adams et al., 2000; Baldwin et al., 2012), prosimian primates (Purushothaman et al., 2012), as well as mice (Scholl et al., 2021; Takahashi, 1985; Tohmi et al., 2014). Furthermore, the identified thalamus-cerebellar connection was supported by a prior finding that the posterior thalamus was connected to inferior cerebellum (Crus I) in individuals with early-onset schizophrenia (Zhang et al., 2021). Importantly, the identified reduced anti-correlation (i.e., hyperconnectivity) between higher order thalamic nuclei and the visual network in SZ aligned well with the findings that SZ patients presented decreased anti-correlation between visual cognition network and the dorsomedial part of thalamus (Gong et al., 2019), as well as the entire thalamus (Ferri et al., 2018), compared to HC. Aberrant FC between the left thalamus and the visual cortex has been documented to be related to attention deficits in schizophrenia (Yamamoto et al., 2018).

Our findings extend prior ROI-based studies by identifying cortico–thalamo–cerebellar circuits in a fully data-driven manner, allowing for subdomain-specific characterization of functional interactions. By integrating information across multiple brain domains, this approach captures coordinated patterns of variation that may be overlooked when examining individual regions or connections in isolation. The consistent replication of this circuit across independent cohorts (COBRE and MPRC) further supports its robustness and potential relevance as a circuit-level marker of cognitive dysfunction in schizophrenia.

The approach and findings presented in this study should be considered in context with its strengths and limitations. The aNy-way ICA framework flexibly integrates multidomain fMRI data while allowing distinct spatial scales across domains, facilitating finer-grained exploration of brain circuits underlying complex brain disorders. One potential limitation is that aNy-way ICA optimizes multimodal linkage using IVA-G within each minibatch, which may limit scalability to very large datasets due to the inherent time complexity of the IVA-G model (Anderson et al., 2012; Sun et al., 2023; Vu et al., 2024). Future work may address this limitation by incorporating more efficient variants of IVA models (Gabrielson et al., 2023; Sun et al., 2023; Vu et al., 2024). Additionally, while this study focuses on identifying circuit-level alterations, further investigation is needed to examine how these patterns are modulated by clinical heterogeneity, symptom profiles, and treatment effects.

In conclusion, we identified and replicated a cortico–thalamo–cerebellar functional circuit associated with cognitive deficits in schizophrenia. This circuit highlights coordinated functional linkages among higher-order thalamic nuclei, cortical networks, and cerebellar regions, and its alteration differentiates individuals with schizophrenia from healthy controls and relates to multiple domains of cognitive impairment. These findings underscore the clinical relevance of cortico–thalamo–cerebellar connectivity and provide insight into circuit-level mechanisms underlying cognitive dysfunction in schizophrenia.

## Acknowledgement

We greatly appreciate individuals who participated in this research study. We would also like to thank research coordinators and principal investigators involved in data collection and data sharing. This work is supported in part by NSF grants 2112455 and 2316421 to VDC. This work was also supported by the National Institutes of Health grants: 1U24 RR021992 (Function Biomedical Informatics Research Network) and 1U24 RR025736-01 (Biomedical Informatics Research Network Coordinating Center), and 1R01MH130595 to JL.

## Data and Code Availability

The code for aNy-way ICA is publicly available in the Fusion ICA Toolbox (FIT, https://trendscenter.org/software/FIT/). COBRE data can be downloaded from COINS (https://coins.trendscenter.org/). The FBIRN and MPRC datasets are not fully publicly available due to IRB restrictions; however, access may be granted upon reasonable request to the principal investigators, subject to institutional approval and applicable data use agreements.

## CRediT authorship contribution statement

**KuaiKuai Duan**: Conceptualization, Data curation, Formal analysis, Investigation, Methodology, Project administration, Software, Validation, Visualization, Writing – original draft, Writing – review and editing. **Rogers F. Silva**: Methodology, Writing – review and editing. **Md Abdur Rahaman**: Data curation, Writing – review and editing. **Zening Fu**: Data curation, Writing – review and editing. **Jingyu Liu**: Resources, Writing – review and editing. **Peter Kochunov**: Resources, Writing – review and editing. **Theo G.M. van Erp**: Resources, Writing – review and editing. **Sarah Shultz**: Writing – review and editing. **Vince D. Calhoun**: Conceptualization, Funding acquisition, Resources, Supervision, Visualization, Writing – review and editing.

## Declaration of competing interest

The authors declare that they have no known competing financial interests or personal relationships that could have appeared to influence the work reported in this paper.

## Notes

### Competing Interest Statement

The authors have declared no competing interest.

### Summary of Updates

Revised the Abstract, Introduction, Discussion, and Conclusion to emphasize biological interpretation, clinical relevance, replication, and cognitive associations. Removed the simulation section and related figures/results because the methodological validation was already published previously in the IEEE EMBC paper. Added brief clarification that methodological validation has been established previously and that the current manuscript focuses on application to human neuroimaging data.

