## Supplementary material for "Cortico–thalamo–cerebellar circuit disruptions underlying cognitive deficits in schizophrenia: a multi-cohort multidomain fusion study"

**Supporting information**

**Resting state fMRI data acquisition**

FBIRN: Resting-state fMRI images were collected at seven sites across the United States. Six sites used the 3T Siemens Tim Trio System, and one site used the 3T General Electric Discovery MR750 scanner. All resting-state fMRI images were acquired using a standard gradient-echo planar imaging paradigm: repetition time (TR) = 2 s, echo time (TE) = 30 ms, field of view (FOV) = 220 × 220 mm (64 × 64 matrix), flip angle = 77°, slice thickness = 4 mm, voxel size = 3.44 mm × 3.44 mm × 4 mm, 162 volumes.

COBRE: Resting-state fMRI images were collected on a 3T Siemens Trio scanner with a 12-channel radio frequency coil. All resting-state fMRI data were acquired using a gradient-echo planar imaging sequence with TR = 2 s, TE = 29 ms, flip angle = 75°, slice thickness = 3.5 mm, slice gap = 1.05 mm, field of view 240 mm, matrix size = 64 × 64, voxel size = 3.75 mm × 3.75 mm × 4.55 mm, 150 volumes.

MPRC: Resting-state fMRI data were aggregated from three cohorts scanned using three 3 T Siemens scanners. All fMRI data were collected using a single‐shot gradient‐recalled, echo‐planar imaging (EPI) pulse sequence with the following acquisition parameters: Cohort A: TR = 2000 ms, TE = 27 ms, matrix size = 64 × 64, voxel size = 3.44 mm × 3.44 mm × 4 mm, 150 volumes; Cohort B: TR = 2210 ms, TE = 27 ms, matrix size = 64 × 64, voxel size = 3.44 mm × 3.44 mm × 3.99 mm, 140 volumes; Cohort C: TR = 2000 ms, TE = 30 ms, matrix size = 128 × 128, voxel size = 1.72 mm × 1.72 mm × 4 mm, 444 volumes.

**Supplementary figures**

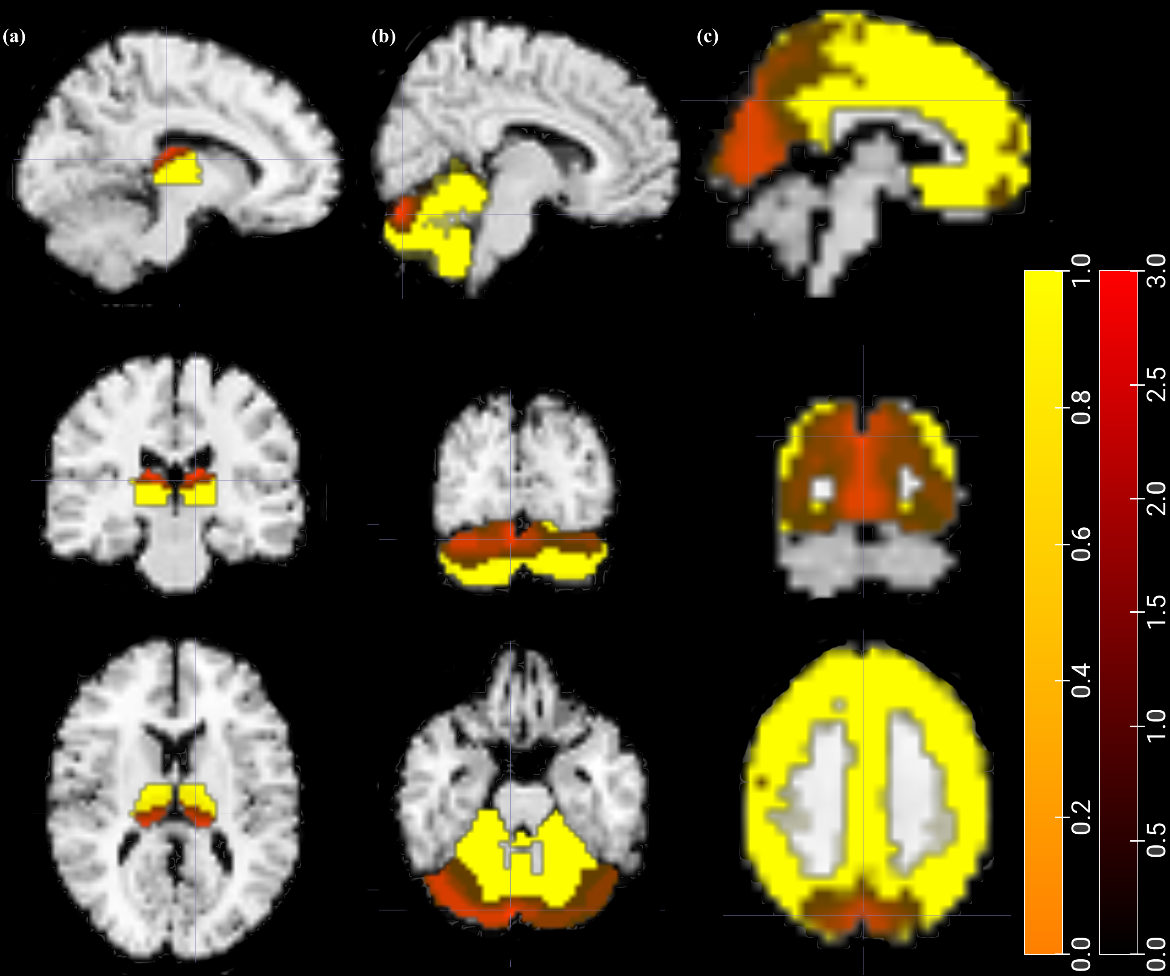

Supplementary Figure 1, The identified (a) thalamus, (b) cerebellum, and (c) cortex component (in red) in SCV6 overlaid to their corresponding masks (in yellow).

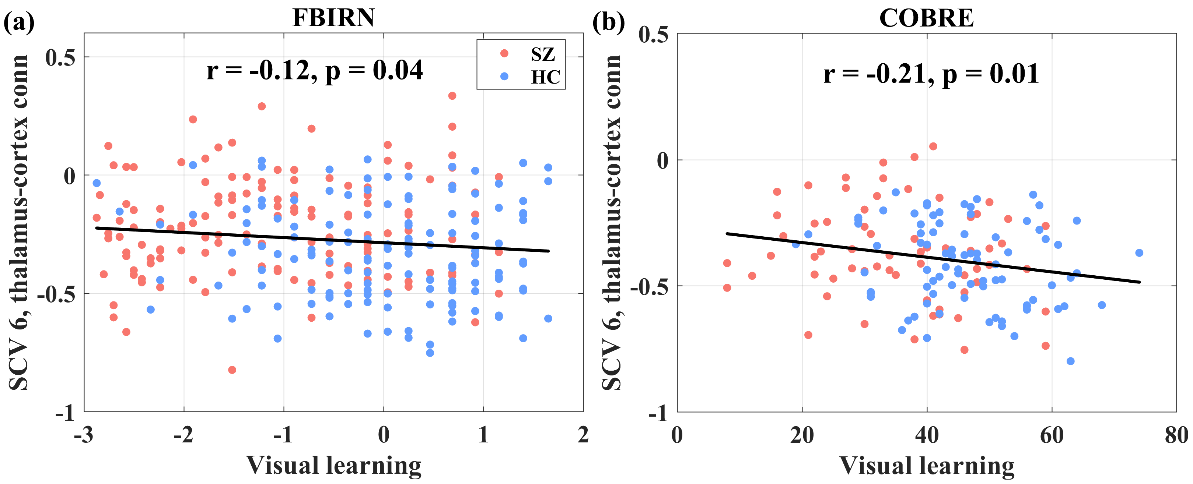

Supplementary Figure 2.  The association between thalamus-cortex connection in SCV 6 and visual learning abilities in the (a) FBIRN (discovery) and (b) COBRE (replication) datasets. Red dots denote individuals with schizophrenia (SZ) and blue dots represent HCs.

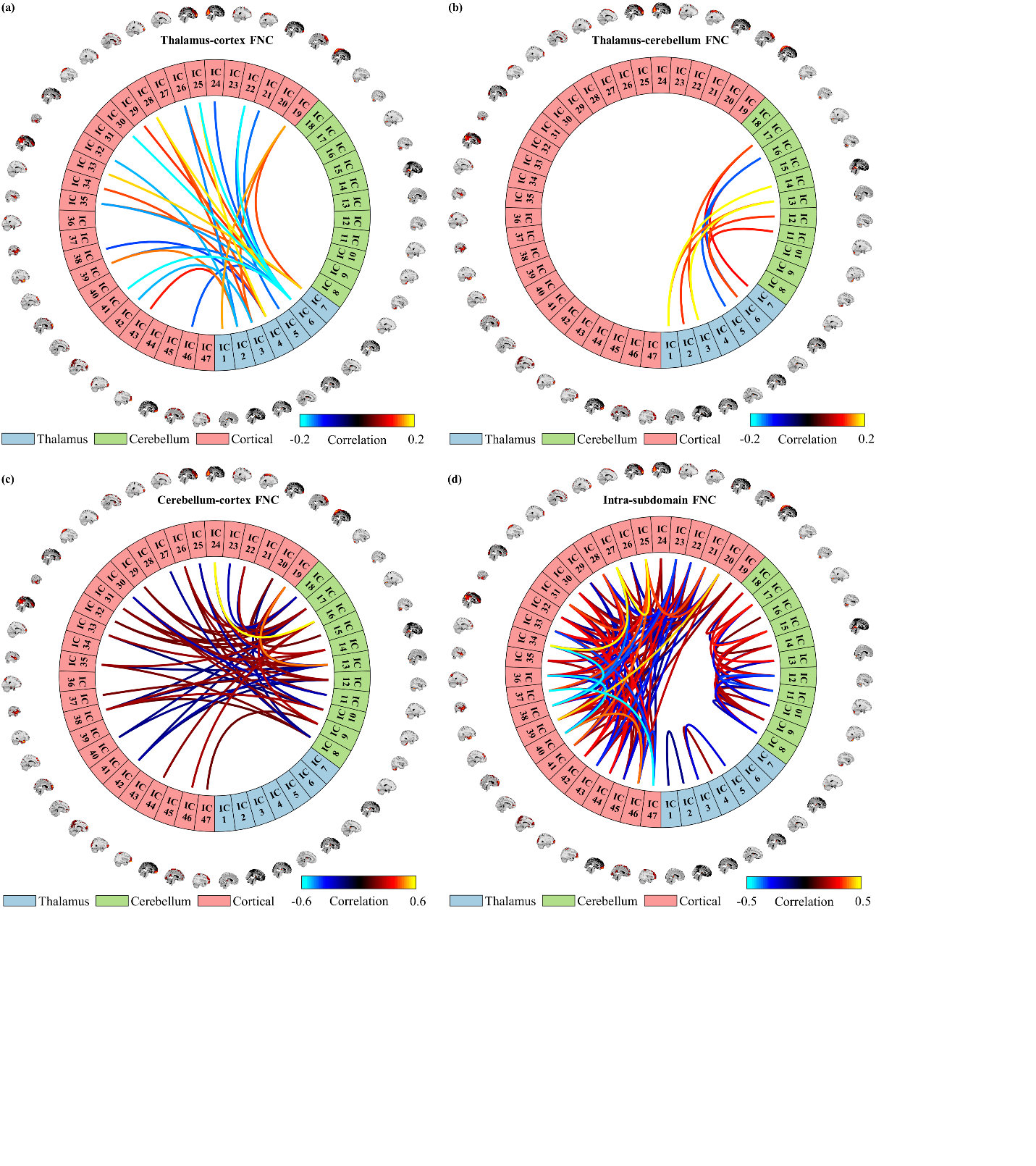

Supplementary Figure 3. The connectogram of the significantly linked whole-brain FNC passing FDR at *p* < 0.05 correction. The connectogram of significant (a) thalamus-cortex FNC, (b) thalamus-cerebellum FNC, (c) cerebellum-cortex FNC, and (d) intra-subdomain (i.e., thalamus, cerebellum, and cortex) FNC. 254 out of 1081 total connection pairs were significantly linked. Among the 254 significant connections, 28 were between thalamus and cortex, 11 were between thalamus and cerebellum, 55 were between cerebellum and cortex, and 160 were intra-subdomain (thalamus/cerebellum/cortex).

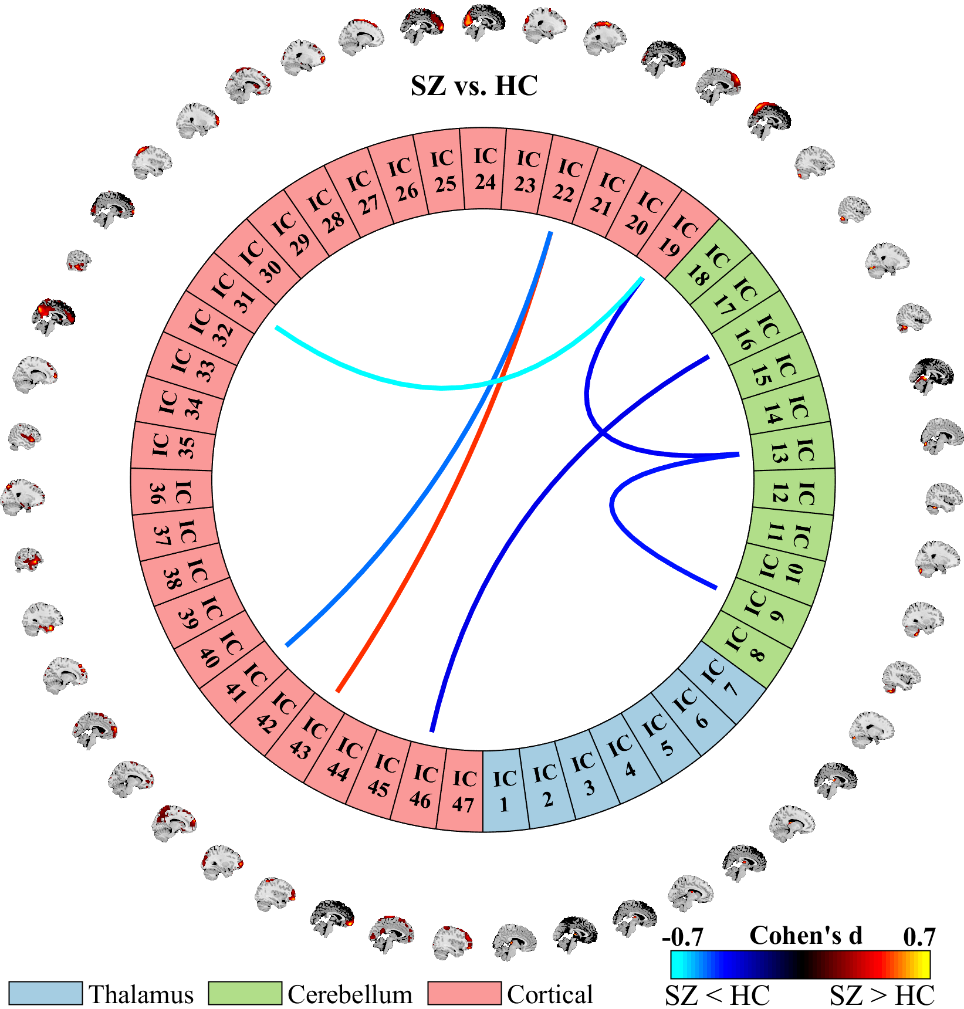

Supplementary Figure 4. The connectogram of FNCs replicated for SZ vs. HC difference in the COBRE dataset (the color of the edge represents Cohen’s d values of SZ vs. HC difference)
